# Semantic-Guided Spatial Representation Learning for Spatial Domain Identification

**DOI:** 10.64898/2026.01.28.700742

**Authors:** Yawen Lu, Yucheng Xu, Junning Feng, Yanlin Zhang

## Abstract

**Motivation:** Spatial domain identification is a fundamental task in spatial transcriptomics analysis, aiming to partition tissue sections into coherent regions that reflect underlying biological organization. Most existing methods rely on gene expression similarity and spatial proximity, which can be insufficient when expression measurements are sparse, noisy, or weakly discriminative. In such settings, representation learning faces ambiguity in determining how spatial neighborhood information and expression-based similarity should be jointly utilized.

**Results:** We present GreS, a spatial domain identification framework that incorporates gene-level semantic priors into spatial representation learning. GreS models spatial adjacency and expression-driven similarity using two complementary neighborhood graphs and leverages aggregated gene semantic information to guide how these views are weighted at the spot level. Rather than redefining neighborhood relationships, semantic priors act as contextual signals that modulate the relative contribution of spatial and expression-based cues during domain inference. We evaluate GreS on diverse spatial transcriptomics datasets, including layered brain tissue, embryonic development, and heterogeneous tumor microenvironments. Across these settings, GreS consistently identifies spatial domains that are structurally coherent and biologically interpretable, outperforming existing methods in quantitative accuracy and qualitative spatial organization. Our ablation analyses further demonstrate that performance gains arise from biologically meaningful and properly aligned semantic information, rather than from increased model complexity or auxiliary features.

**Availability and Implementation:** https://github.com/ai4nucleome/GreS

**Supplementary Information:** A Supplementary file is submitted together with this manuscript.

## Introduction

Spatially resolved transcriptomics (ST) enables the measurement of gene expression while preserving the spatial organization of tissues, providing a powerful framework for studying tissue architecture and microenvironmental heterogeneity. Accurate spatial domain identification is a fundamental yet challenging problem in spatial transcriptomics analysis, aiming to partition tissue sections into coherent regions corresponding to distinct molecular or functional states. Reliable identification of spatial domains underpins many downstream analyses, including tissue organization characterization, spatially variable gene analysis, and the interpretation of disease-associated microenvironments [Rao et al., 2021].

A large number of computational methods have been proposed for spatial domain detection. Early approaches, such as Seurat [Satija et al., 2015] and Scanpy [Wolf et al., 2018], primarily adapt clustering strategies developed for single-cell RNA-seq analysis, applying them directly to ST data with little or no explicit modeling of spatial structure. Subsequent methods, including BayesSpace [Zhao et al., 2021] and Giotto [Chen et al., 2023], incorporate spatial information by imposing spatial smoothness or probabilistic constraints. More recent approaches, like STAGATE [Dong and Zhang, 2022] and GraphST [Long et al., 2023], explicitly construct graphs over tissue locations and employ graph-based or graph neural network models to jointly model gene expression and spatial adjacency, with some methods further integrating histological images. Although these advances have improved performance in well-structured tissues, most existing methods rely predominantly on measured gene expression signals, which are often sparse, noisy, and limited in coverage. Consequently, spatial domains characterized by subtle transcriptional differences or gradual transitions remain difficult to resolve, particularly when expression information alone provides insufficient discriminative power.

One common strategy to mitigate this limitation is to introduce additional sources of information beyond gene expression. For example, several recent methods, such as stLearn [Pham et al., 2023] and SpaGCN [Hu et al., 2021], incorporate histological images to provide complementary morphological cues that help stabilize spatial domain inference. These approaches demonstrate that augmenting ST data with external, orthogonal information can substantially improve domain detection when expression signals are weak or ambiguous. However, histological images are not always available, and their integration introduces additional technical challenges related to image preprocessing, feature extraction, and spatial alignment.

An alternative and largely unexplored source of complementary information lies in the biological semantics of genes themselves. Most existing methods treat genes as independent numerical features, ignoring their functional meaning and regulatory relationships. However, spatial domains are often defined by coordinated activity of functionally related gene programs rather than by large expression changes in individual genes. Recent work in single-cell and related transcriptomic analyses, such as GenePT [Chen and Zou, 2025], has shown that large language models can encode gene functional information from curated descriptions and biological knowledge sources, providing semantic representations that serve as informative biological priors. Incorporating such gene semantic information has the potential to enrich spot-level representations and compensate for sparse or noisy expression measurements. Nevertheless, its use in spatial transcriptomics, and in particular for spatial domain detection, remains limited. Moreover, effectively integrating gene semantics into spatial models requires addressing additional challenges, including interdependent gene functions and the need to balance semantic and spatial cues across heterogeneous tissue regions, motivating a task-specific integration strategy rather than a fixed fusion scheme.

Here, we propose GreS, a semantic-guided framework for spatial domain identification that leverages gene functional knowledge as a biologically informed prior. GreS incorporates gene-level semantic information derived from curated functional descriptions and integrates it with spatial transcriptomics data to guide representation learning beyond expression patterns alone. By adaptively balancing spatial proximity and functional similarity, GreS enables more robust identification of spatial domains across heterogeneous tissue contexts. Extensive evaluations on brain, developmental, and tumor datasets demonstrate that GreS consistently improves domain coherence and biological interpretability compared with existing methods.

## Materials and methods

### Model architecture overview

GreS takes spatial transcriptomics data as input and learns spot-level representations through semantic-guided spatial modeling. As shown in Fig. 1a, the model starts from the observed gene expression, where each spatial spot is characterized by its expressed genes and corresponding expression values. Based on these inputs, GreS first constructs spot-level semantic descriptors (Fig. 1b) by encoding gene descriptions, refining them through a gene regulatory network, and aggregating the refined gene semantics according to each spot’s expression profile. These semantic descriptors provide a functional context for downstream representation learning. Next, GreS builds two complementary graphs over spatial spots (Fig. 1c): a feature graph capturing functional similarity between spots and a spatial graph capturing physical neighborhood relationships. Separate graph convolutional encoders are applied to obtain feature-view and spatial-view embeddings. Finally, as illustrated in Fig. 1d, GreS uses the spot-level semantic descriptor to adaptively gate and fuse the two view-specific embeddings into a unified representation. GreS is trained in an autoencoding framework, where the fused spot representations are optimized to reconstruct the input gene expression via a zero-inflated negative binomial decoder, together with additional regularization objectives that preserve spatial consistency and cross-view complementarity. After training, the learned spot embeddings are extracted and used as input to K-means algorithm to identify spatial domains.

**Fig. 1:**
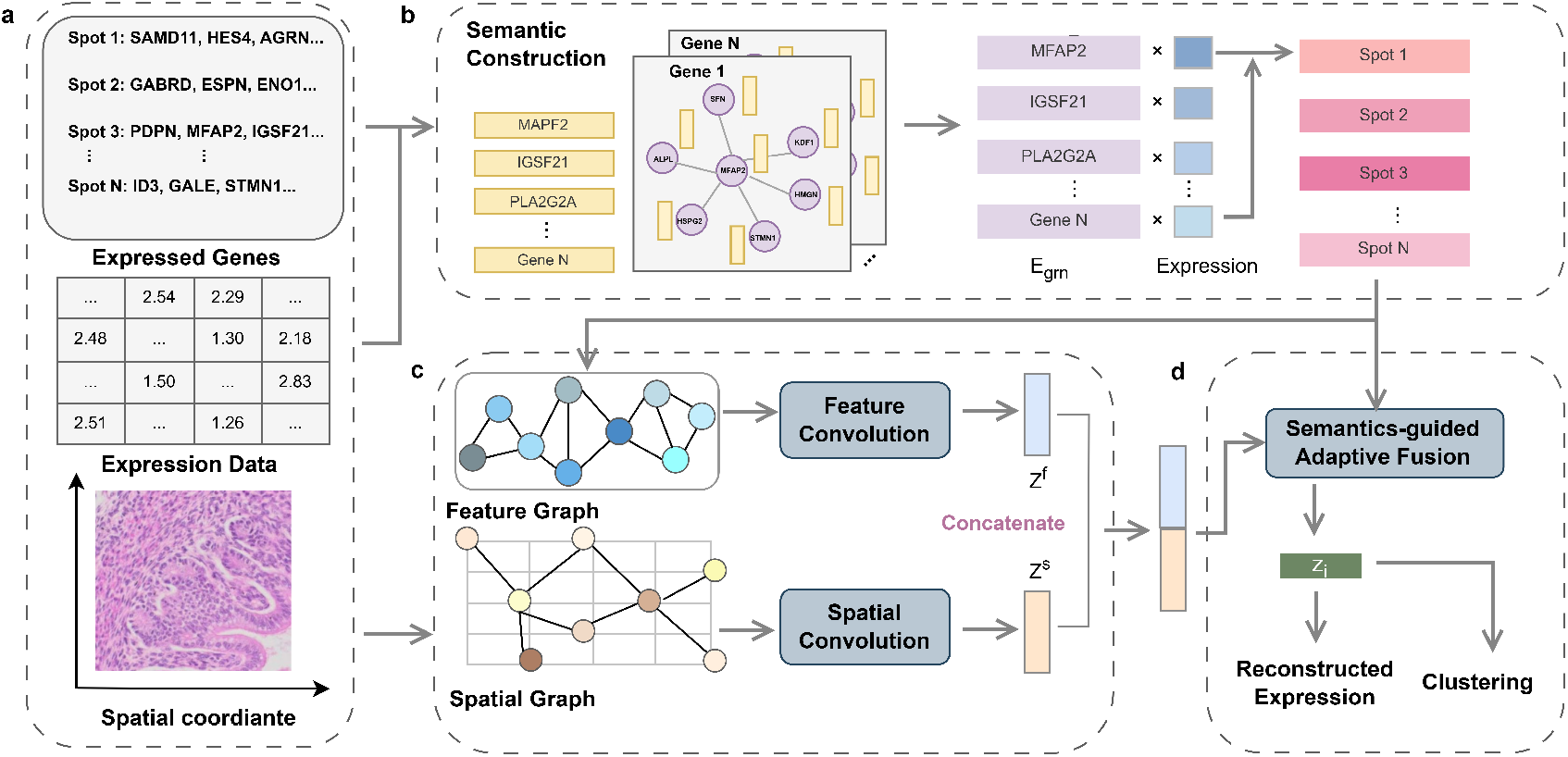
Overview of GreS. (a) Input spatial transcriptomics data, where each spot is represented by its expressed genes and expression values. (b) Semantic construction: gene functional descriptions are encoded, refined via gene regulatory network (GRN) propagation, and aggregated by expression to obtain spot-level semantic descriptors. (c) Dual-view graph encoding: a feature graph and a spatial graph are constructed over spots and processed by graph convolutions to produce feature-view embeddings *Z*^*f*^ and spatial-view embeddings *Z*^*s*^. Semantics-guided adaptive fusion: spot-level semantic descriptors are used to gate and fuse *Z*^*f*^ and *Z*^*s*^ into the final representation *z*_*i*_ for spatial domain identification.

### GRN-guided semantic construction

GreS introduces a semantic construction module to incorporate gene functional knowledge as a biologically informed prior for spatial representation learning (Fig. 1b). The goal of this module is to summarize the functional context of genes expressed at each spatial spot, producing a compact semantic descriptor that can guide downstream graph-based modeling. This module operates in three stages: (i) encoding gene functional descriptions into semantic embeddings, (ii) refining these embeddings using gene regulatory networks, and (iii) aggregating refined gene semantics into spot-level semantic descriptors based on expression.

### Gene-level semantic embeddings

For each gene *g*, we retrieve a curated textual summary describing its known biological functions from the NCBI Gene database [Brown et al., 2015]. These descriptions are encoded into semantic vectors using `text-embedding-3-large`, resulting in a gene semantic embedding 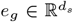 . Stacking embeddings for the *G* selected genes yields a gene semantic matrix

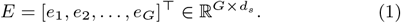

Not all genes have available functional descriptions. For genes without curated annotations, we simply set their semantic embeddings to zero vectors.

### GRN-based refinement of gene semantics

Gene functions are not independent but organized through regulatory and signaling relationships. To incorporate this structure, we refine gene semantic embeddings using GRNs derived from the NicheNet prior [Browaeys et al., 2023]. The GRNs integrates curated ligand–receptor interactions, intracellular signaling pathways, and transcriptional regulation, providing a biologically grounded dependency structure among genes.

After restricting the GRN to the selected genes, we obtain a weighted adjacency matrix *A*_grn_ ∈ ℝ^*G×G*^, which is row-normalized to *Â*_grn_. Gene semantic embeddings are refined through one-step graph propagation:

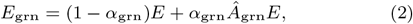

where *α*_grn_ is a hyperparameter, and we set *α*_grn_ = 0.2 in all experiments.

This operation propagates functional information along biologically meaningful regulatory relationships, producing refined gene semantics that jointly encode functional meaning and regulatory context.

### Expression-weighted spot-level semantic descriptors

Spatial domain identification operates at the level of spatial spots rather than individual genes. We therefore aggregate refined gene semantic embeddings into a spot-level semantic descriptor based on gene expression.

For each spot *i*, we compute a semantic descriptor *u*_*i*_ as an expression-weighted average of refined gene semantics:

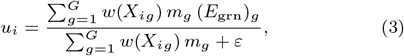

where *X*_*ig*_ denotes the expression of gene *g* at spot *i, w*(·) is a non-negative weighting function proportional to expression (default *w*(*x*) = *x*), and *ε* is a small constant for numerical stability.

The resulting matrix

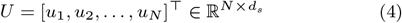

provides a compact semantic summary for each spatial spot.

### Dual-view graph construction and encoding

After obtaining the spot semantic descriptors *U*, GreS learns two complementary spot representations by encoding the tissue under two neighborhood definitions (Fig. 1c): (i) a *spatial view* that captures local physical continuity, and (ii) a *feature view* that captures functional similarity informed by gene semantics.

#### Graph construction

We construct two graphs over the same set of *N* spots. The spatial graph *A*_*s*_ is built from spatial coordinates *P* by connecting each spot to its *k* nearest neighbors under Euclidean distance:

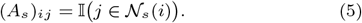

The feature graph *A*_*f*_ is constructed in the semantic space using cosine similarity between spot descriptors {*u*_*i*_}:

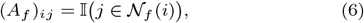

where 𝒩_*f*_ (*i*) denotes the top-*k* most similar spots to *i* measured by cosine similarity on {*u*_*i*_}. We use the same *k* for both graphs.

#### Dual-view graph encoding

Both graphs share the same node features given by the expression matrix *X*, but differ in their connectivity. For each view *v* ∈ {*s, f* }, we add self-loops and apply row normalization:

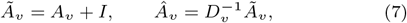

where *D*_*v*_ is the degree matrix of Ã_*v*_ . We then employ two independent two-layer GCN encoders to obtain view-specific embeddings, and set the hidden dimension to 256 and output dimension to 128 in GCN.

The resulting embeddings *Z*^*s*^ and *Z*^*f*^ provide complementary characterizations of tissue organization: *Z*^*s*^ emphasizes spatial smoothness induced by physical adjacency, while *Z*^*f*^ captures functionally similar regions connected through semantic similarity. These two representations are subsequently integrated by the semantics-guided adaptive fusion module.

### Semantics-guided adaptive fusion

GreS yields two view-specific embeddings for each spatial spot: a spatial-view embedding 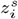 capturing physical neighborhood continuity and a feature-view embedding 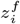 capturing functional similarity. We regard these two embeddings as alternative explanations of the local tissue state. Since neither view is uniformly reliable across all tissue regions, the objective of the fusion module is to determine how much each view should contribute for a given spot.

### Semantic-conditioned gating

We first concatenate the two view embeddings to form the gate input:

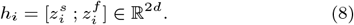

The gating network uses this joint representation to assess the relative reliability of the two views. To incorporate biological context into this decision, the spot semantic descriptor *u*_*i*_ is used to modulate the gate input through a feature-wise affine transformation:

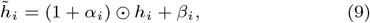

where the modulation parameters *α*_*i*_ and *β*_*i*_ are predicted from *u*_*i*_. This step allows semantic information to influence how spatial and functional cues are evaluated, rather than being directly fused as a third representation.

### Channel-wise fusion

From the modulated representation 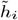, a channel-wise gate *g*_*i*_ ∈ (0, 1)^*d*^ is predicted by a gating network. The gate specifies, for each feature channel, the relative contribution of the spatial and feature views. The final fused embedding is obtained as follows:

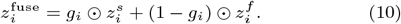

A lightweight post-fusion projection is applied to obtain the final representation *z*_*i*_, which is used for downstream reconstruction and spatial domain identification.

### Learning objectives and optimization

The deep learning part of GreS is trained end-to-end by optimizing a weighted sum of three objectives: (i) ZINB-based expression reconstruction, (ii) spatial neighborhood regularization, and (iii) cross-view decorrelation:

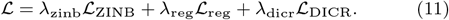

The loss weights *λ*_zinb_, *λ*_reg_, and *λ*_dicr_ control the relative contributions of the three objectives. We selected these hyperparameters via a grid search on a representative validation slice, with each weight varied in the range [0.1, 1]. The configuration that achieved the best clustering performance on the validation slice was *λ*_zinb_ = 1.0, *λ*_reg_ = 0.1, and *λ*_dicr_ = 0.1. This set of hyperparameters was then fixed and applied to all remaining slices and datasets. We additionally experimented with re-tuning the loss weights on other slices, but found that this configuration consistently yielded competitive performance.

### ZINB reconstruction loss

Following common practice for sparse count data, we decode *Z* to ZINB parameters (i.e, *π, θ*, and *µ*) and minimize the negative log-likelihood of the observed expression matrix *X*:

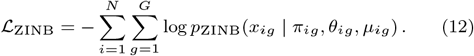

### Spatial neighborhood regularization

To encourage local spatial smoothness without collapse, we sample positive pairs ℰ ^+^ from edges of the spatial graph *A*_*s*_ and negative pairs ℰ^−^ from non-edges. Let *p*_*ij*_ = *σ*(sim(*z*_*i*_, *z*_*j*_ )) where sim(·, ·) is cosine similarity (on ℓ_2_-normalized embeddings). We optimize a binary cross-entropy objective:

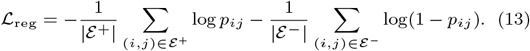

### Dual-view decorrelation loss

To promote complementary information between the spatial and feature views, we penalize cross-view correlations between channels of *Z*^*s*^ and *Z*^*f*^ . Let *S* = norm(*Z*^*s*^)^⊤^norm(*Z*^*f*^ ) ∈ ℝ^*d×d*^, where norm(·) denotes ℓ_2_ normalization across spots (row-wise on *Z*). We minimize deviation from the identity:

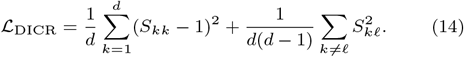

We optimize ℒusing Adam; after training, the learned embedding *Z* is used for downstream clustering to obtain spatial domains.

### Spatial domain detection

For a given spatial transcriptomics dataset, GreS is first trained end-to-end using the semantic-guided spatial representation learning framework described above. Through this training process, the model learns a low-dimensional embedding *Z* for each spatial spot. After training converges, the learned spot embeddings *Z* are extracted and used as input to a clustering algorithm. In this work, we apply *k*-means clustering to *Z* to partition spatial spots into distinct domains, with the number of clusters set according to the available ground-truth annotations for benchmarking or a user defined value in real application.

### Datasets, preprocessing, and evaluation protocol

#### Spatial transcriptomics datasets

We evaluated GreS using three publicly available spatial transcriptomics datasets spanning distinct biological contexts (Supplementary Table S1). The human dorsolateral prefrontal cortex (DLPFC) dataset was generated using 10x Visium and consists of 12 tissue sections from three donors with manually annotated cortical layers [Maynard et al., 2021]. The mouse embryo (ME) dataset was acquired using Stereo-seq and comprises five E9.5 embryo slices annotated into 13 developmental regions [Chen et al., 2022, Liu et al., 2020]. The human breast cancer (HBC) dataset is a 10x Visium sample representing a heterogeneous tumor microenvironment and is manually annotated into 20 spatial domains [Xu et al., 2024].

### Data preprocessing

All datasets were processed using a standardized pipeline implemented with Scanpy [Wolf et al., 2018]. Genes expressed in fewer than 100 spots were filtered out, after which the top *N*_HVG_ highly variable genes were selected using the Seurat v3 dispersion-based method [Satija et al., 2015], with *N*_HVG_ set to 3000 by default. For model input, gene expression counts were normalized to a total count of 10^4^ per spot to account for sequencing depth, followed by log-transformation and scaling to zero mean and unit variance with values clipped at 10. Raw integer count matrices were retained to supervise the ZINB reconstruction loss. Spatial coordinates were used directly to construct spatial neighborhood graphs.

### Baseline methods

We compared GreS against nine representative spatial domain identification methods spanning diverse methodological paradigms, including contrastive learning–based approaches (stDCL [Yu et al., 2025], GraphST [Long et al., 2023], and GAAEST [Wang et al., 2024]), graph neural network–based models (STAGATE [Dong and Zhang, 2022], Spatial-MGCN [Wang et al., 2023], SEDR [Xu et al., 2024], and SpaGCN [Hu et al., 2021]), a statistical method (stLearn [Pham et al., 2023]), and a semantic-based baseline (SemST [Long et al., 2025]). All baseline methods were executed using their official implementations with recommended parameter settings. Clustering was performed in an unsupervised manner, with the number of clusters fixed to match the ground-truth annotations for each dataset, following common benchmarking practice.

### Evaluation metrics

Clustering performance was assessed using four standard metrics: Adjusted Rand Index (ARI), Normalized Mutual Information (NMI), Clustering Accuracy (ACC), and Fowlkes–Mallows Index (FMI).

## Results

To comprehensively evaluate the effectiveness of GreS, we conducted a series of benchmarking experiments on three publicly available spatial transcriptomics datasets representing distinct biological contexts (see Methods). These datasets encompass well-organized layered tissue, complex developmental structures, and heterogeneous pathological microenvironments. We compared GreS with nine representative spatial domain identification methods in terms of clustering accuracy, spatial coherence, and biological interpretability. In addition, we performed ablation studies to assess the specific contributions of the semantic-guided fusion mechanism.

### GreS accurately identifies laminar spatial domains in brain tissue

We first examined whether GreS can accurately identify laminar spatial domains in layered brain tissue using the DLPFC dataset, a widely adopted benchmark with high-quality manual annotations of cortical layers. Across all 12 tissue sections from three donors, GreS demonstrated consistently strong clustering performance. As shown in Figure 2a, GreS achieved the highest median ARI among all compared methods and showed a distribution shifted toward higher ARI values, indicating stable performance across donors and slices.

**Fig. 2:**
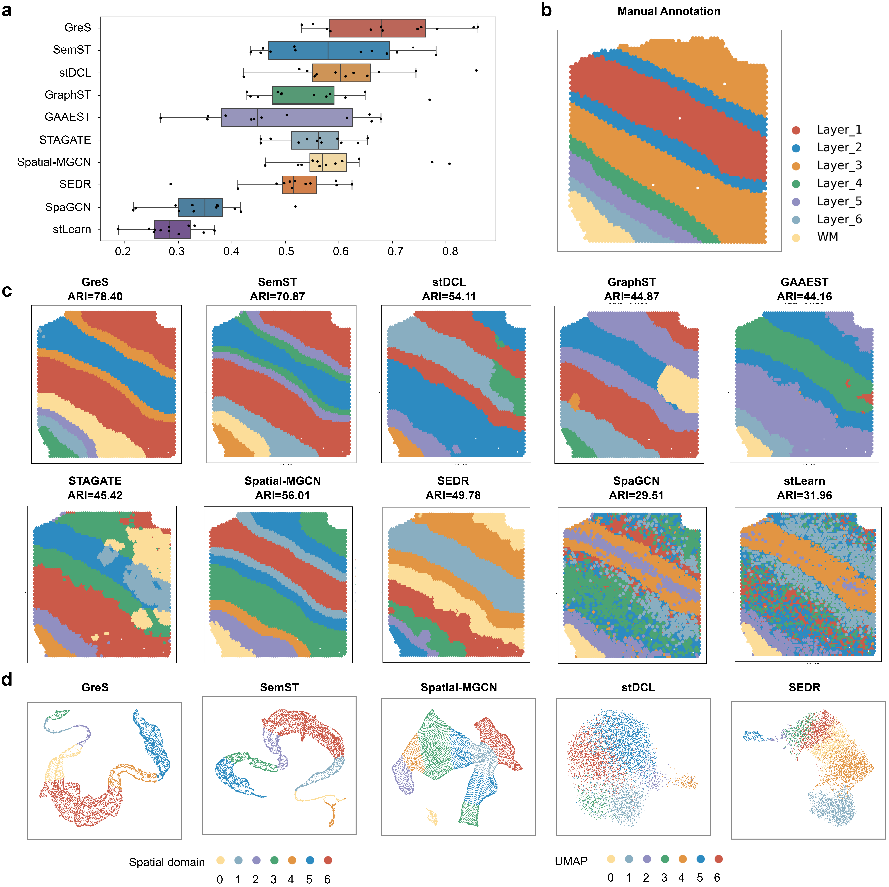
Analysis of the DLPFC dataset: (a) Boxplot of ARI scores across all 12 DLPFC slices for ten methods; (b) Manually annotated domains for slice 151509; (c) Domains predicted by GreS and alternative methods on slice 151509; (d) UMAP plots of the latent representations learned by selected methods for slice 151509.

Detailed quantitative comparisons for three representative sections from different donors are reported in Table 1. Across these sections, GreS exhibited consistently strong performance across all four evaluation metrics (ARI, NMI, ACC, and FMI), distinguishing itself from both semantic-based and graph-based alternatives in terms of overall ranking stability across donors. On section 151509, most graph-based methods achieved moderate performance, with noticeable variation across metrics. In contrast, GreS achieved uniformly high scores across all four metrics, indicating more coherent recovery of spatial domain structure compared with other methods. On section 151672, several leading methods, including SemST, Spatial-MGCN, and GraphST, also performed well, reflecting a generally strong signal in this slice. In this setting, GreS remained competitive and achieved the highest overall scores, demonstrating that its advantage is maintained even when multiple methods perform favorably. On section 151676, several existing methods exhibited pronounced performance variability across metrics, whereas GreS maintained stable performance on ARI, NMI, ACC, and FMI.

**Table 1.**
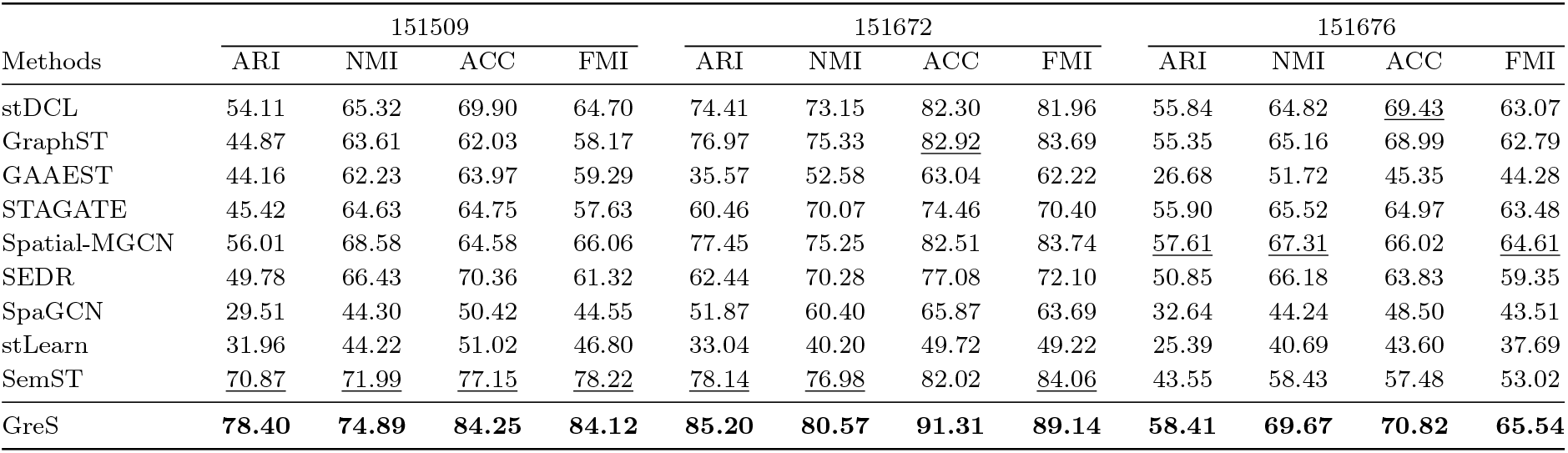
Quantitative comparison on DLPFC slices 151509, 151672, and 151676. ARI, NMI, ACC, and FMI are reported as percentages (values in [0, 1] multiplied by 100). The best and second best results are highlighted in **bold** and underlined, respectively.

| Methods | 151509 |  |  |  | 151672 |  |  |  | 151676 |  |  |  |
| --- | --- | --- | --- | --- | --- | --- | --- | --- | --- | --- | --- | --- |
|  | ARI | NMI | ACC | FMI | ARI | NMI | ACC | FMI | ARI | NMI | ACC | FMI |
| stDCL | 54.11 | 65.32 | 69.90 | 64.70 | 74.41 | 73.15 | 82.30 | 81.96 | 55.84 | 64.82 | <u>69.43</u> | 63.07 |
| GraphST | 44.87 | 63.61 | 62.03 | 58.17 | 76.97 | 75.33 | <u>82.92</u> | 83.69 | 55.35 | 65.16 | 68.99 | 62.79 |
| GAAEST | 44.16 | 62.23 | 63.97 | 59.29 | 35.57 | 52.58 | 63.04 | 62.22 | 26.68 | 51.72 | 45.35 | 44.28 |
| STAGATE | 45.42 | 64.63 | 64.75 | 57.63 | 60.46 | 70.07 | 74.46 | 70.40 | 55.90 | 65.52 | 64.97 | 63.48 |
| Spatial-MGCN | 56.01 | 68.58 | 64.58 | 66.06 | 77.45 | 75.25 | 82.51 | 83.74 | <u>57.61</u> | <u>67.31</u> | 66.02 | <u>64.61</u> |
| SEDR | 49.78 | 66.43 | 70.36 | 61.32 | 62.44 | 70.28 | 77.08 | 72.10 | 50.85 | 66.18 | 63.83 | 59.35 |
| SpaGCN | 29.51 | 44.30 | 50.42 | 44.55 | 51.87 | 60.40 | 65.87 | 63.69 | 32.64 | 44.24 | 48.50 | 43.51 |
| stLearn | 31.96 | 44.22 | 51.02 | 46.80 | 33.04 | 40.20 | 49.72 | 49.22 | 25.39 | 40.69 | 43.60 | 37.69 |
| SemST | <u>70.87</u> | <u>71.99</u> | <u>77.15</u> | <u>78.22</u> | <u>78.14</u> | <u>76.98</u> | 82.02 | <u>84.06</u> | 43.55 | 58.43 | 57.48 | 53.02 |
| GreS | **78.40** | **74.89** | **84.25** | **84.12** | **85.20** | **80.57** | **91.31** | **89.14** | **58.41** | **69.67** | **70.82** | **65.54** |

This contrast highlights the ability of GreS to deliver consistent and balanced clustering performance across sections with varying signal characteristics.

Spatial visualizations further illustrate how GreS identifies laminar organization. Figure 2c compares spatial domain assignments on section 151509. The ground-truth annotation reveals a clear laminar structure spanning cortical Layers 1–6 and white matter. GreS successfully reconstructed this structure with smooth, contiguous boundaries that closely match the annotation. In contrast, several baseline methods struggled to distinguish adjacent layers or produced fragmented patterns. For example, STAGATE and GAAEST failed to clearly separate Layers 4 and 5, while SpaGCN and stLearn generated scattered clusters lacking spatial coherence. Although stDCL and SemST recovered the overall laminar organization, GreS achieved sharper boundary delineation, particularly at transitions between deep cortical layers and white matter.

To further assess how GreS captures laminar structure at the representation level, we visualized spot embeddings using UMAP (Figure 2d). GreS embeddings formed a well-ordered trajectory consistent with the intrinsic spatial progression of cortical layers, with distinct clusters corresponding to individual layers. In comparison, embeddings from methods such as stDCL and SEDR exhibited substantial overlap between clusters, reflecting weaker discriminability in the learned feature space. Together, these results indicate that GreS accurately identifies laminar spatial domains in brain tissue by integrating spatial structure with biologically informed representations.

### GreS resolves fine-grained spatial domains during embryonic development

We next evaluated GreS on a high-resolution Stereo-seq mouse embryo dataset (E9.5), which contains five annotated slices (E1S1 and E2S1–E2S4) with 13 developmental regions. Quantitative comparisons are summarized in Table 2 and Figure 3c. GreS achieved the highest average ARI across slices (0.4278). Specifically, GreS obtained the best ARI on four of the five slices (E2S1–E2S4), with the only exception being E1S1, where SemST slightly outperformed GreS (ARI: 0.4013 vs. 0.3956).

**Table 2.** Detailed ARI scores (scaled by 100) on five mouse embryo slices. The best performance is highlighted in **bold**, and the second best is underlined.

| Slice | GreS | SemST | stDCL | GraphST | GAAEST | STAGATE | Spatial-MGCN | SEDR | SpaGCN | stLearn |
| --- | --- | --- | --- | --- | --- | --- | --- | --- | --- | --- |
| E1S1 | <u>39.56</u> | **40.13** | 29.29 | 28.57 | 29.71 | 32.66 | 29.60 | 28.88 | 32.08 | 31.32 |
| E2S1 | **41.78** | 25.56 | 28.91 | 25.04 | 22.06 | <u>31.95</u> | 31.52 | 29.67 | 38.86 | 29.80 |
| E2S2 | **42.02** | 34.85 | 34.69 | 34.26 | 29.78 | 37.05 | <u>41.49</u> | 38.48 | 37.88 | 35.82 |
| E2S3 | **45.15** | 33.55 | 36.34 | 37.10 | 33.77 | 39.60 | 39.18 | <u>43.58</u> | 43.57 | 37.17 |
| E2S4 | **45.38** | 36.29 | 31.65 | 33.99 | 27.20 | 37.23 | 30.21 | 36.16 | 33.90 | <u>41.90</u> |
| **Avg.** | **42.78** | 34.08 | 32.18 | 31.79 | 28.50 | 35.70 | 34.40 | 35.35 | <u>37.26</u> | 35.20 |

**Fig. 3:**
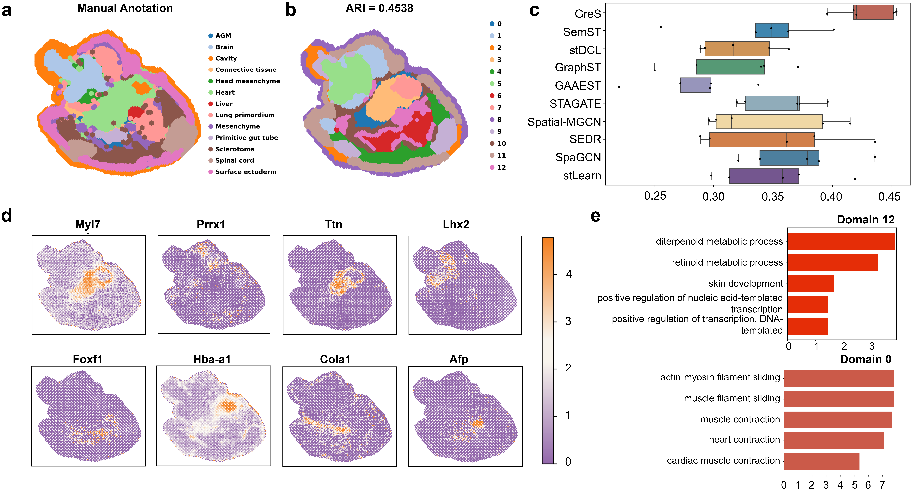
Analysis of the mouse embryo dataset. (a) Manual annotation of slice E2S4. (b) Spatial domains identified by GreS on slice E2S4. (c) Distribution of ARI scores comparing GreS with existing methods across five embryo slices. (d) Spatial expression patterns of representative marker genes. Gene Ontology enrichment analysis for two representative GreS-identified domains, highlighting skin development–related processes (Domain 12) and heart contraction–related processes (Domain 0).

We further examined the spatial domains predicted on slice E2S4 as a representative example (Figure 3a,b), where GreS achieved an ARI of 0.4538. The inferred domains form spatially coherent regions that align well with the manual annotation and preserve fine-grained anatomical structure. In particular, GreS delineates the developing heart region (Cluster 0) and separates it from surrounding tissues (Cluster 8), while also resolving smaller structures that are spatially adjacent to neighboring regions.

The biological validity of the identified domains was confirmed by the spatial expression of canonical marker genes (Figure 3d). The heart-associated domain shows localized expression of contractile genes *Myl7* and *Ttn*, whereas *Lhx2* highlights forebrain regions. Additional markers exhibit spatially restricted patterns consistent with known embryonic anatomy, including *Foxf1* (lung primordium), *Afp* (liver/gut region), and *Hba-a1* (erythroid populations). Structural components are also well-resolved, with *Prrx1* [Zhong et al., 2020] and *Col1a1* [Fullard et al., 2021] delineating mesenchymal and connective tissues.

Finally, Gene Ontology enrichment analysis provides functional support for the identified domains (Figure 3e). The heart-associated cluster (Cluster 0) is enriched for terms related to heart contraction and muscle filament sliding, while another representative cluster (Cluster 12) shows enrichment for skin development and retinoid metabolic processes.

Together, these results indicate that incorporating gene-level semantic information enables GreS to capture spatial domains in embryonic tissues where organization is driven by subtle functional programs.

### GreS delineates heterogeneous spatial domains in tumor microenvironments

We next evaluated GreS on the HBC dataset to assess its ability to delineate heterogeneous spatial domains in a pathological tissue context. Quantitative results are summarized in Table 3. Across the four evaluation metrics, GreS achieved the best overall performance, ranking first in ARI, ACC, and FMI, while remaining competitive in NMI. Although Spatial-MGCN achieved the highest NMI, it exhibited lower ARI and FMI compared with GreS, indicating that high mutual information alone does not necessarily translate into accurate domain assignments. In contrast, GreS consistently performed well across all metrics. This balanced performance suggests that GreS more effectively captures the underlying spatial structure of heterogeneous tumor tissues, rather than optimizing for a single clustering criterion.

**Table 3.** Quantitative comparison on the human breast cancer dataset. ARI, NMI, ACC, and FMI are reported as percentages (values in [0, 1] multiplied by 100). The best performance is highlighted in **bold**, and the second best is underlined.

| Method | ARI | NMI | ACC | FMI |
| --- | --- | --- | --- | --- |
| stDCL | 54.40 | 68.25 | 56.71 | 57.75 |
| GraphST | 56.78 | <u>69.39</u> | 58.00 | 59.96 |
| GAAEST | 51.25 | 67.50 | 58.79 | 55.47 |
| STAGATE | 49.00 | 63.93 | 51.26 | 52.75 |
| Spatial-MGCN | 65.24 | **69.93** | 63.82 | <u>67.83</u> |
| SEDR | 37.04 | 64.46 | 46.26 | 41.64 |
| SpaGCN | 48.16 | 61.17 | 52.79 | 51.94 |
| stLearn | 49.37 | 60.20 | 53.29 | 53.07 |
| SemST | <u>68.44</u> | 67.57 | <u>66.46</u> | 27.50 |
| **GreS** | **69.82** | 68.01 | **66.56** | **72.11** |

We examined the spatial domains inferred by GreS on the human breast cancer dataset through direct comparison with the manual annotation and representative existing methods (Figure 4a). The manual annotation reveals complex and heterogeneous spatial organization, including multiple invasive ductal carcinoma (IDC), ductal carcinoma in situ (DCIS), and tumor–edge regions with irregular and non-contiguous boundaries. The spatial domains identified by GreS form contiguous regions while preserving fine-scale spatial heterogeneity that closely reflects the structure of the manual annotation. Among existing methods, SemST also recovers much of the large-scale tumor structure and produces spatially coherent domains that align well with the manual annotation. Spatial-MGCN and GraphST, while generating spatially continuous domains, frequently aggregate heterogeneous tumor, edge, and stromal regions into larger mixed clusters, resulting in weaker alignment with the annotated histological structure.

**Fig. 4:**
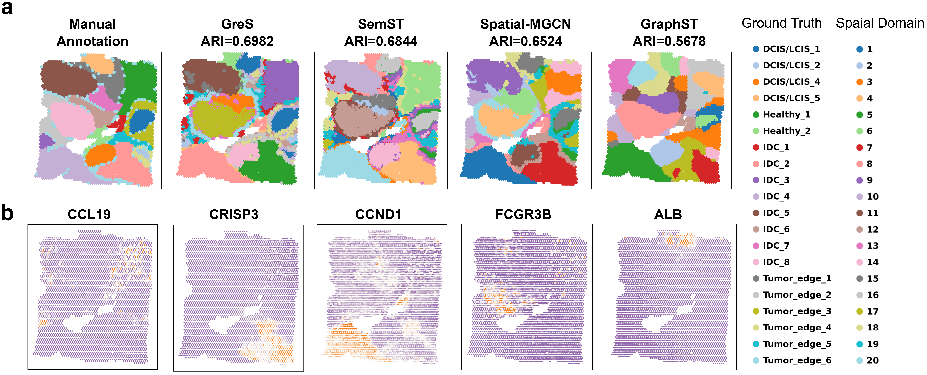
Analysis of the Human Breast Cancer data. (a) Comparison of spatial domains identified by GreS, SemST, Spatial-MGCN, and GraphST against the manual annotation. (b) Spatial expression patterns of key marker genes.

As show in Figure 4b, marker gene analysis further supports the biological plausibility of the identified domains. Proliferative tumor regions showed elevated expression of the cell-cycle regulator *CCND1*, whereas immune-associated markers such as *FCGR3B* and *CCL19* localized to regions consistent with immune infiltration. Furthermore, the secretory protein *CRISP3* [Volpert et al., 2020] exhibited specific localization in ductal structures, and the plasma protein *ALB* [Fanali et al., 2012] clearly demarcated the healthy stromal regions, separating them from the tumor mass. These patterns align with the spatial domains identified by GreS, indicating that the inferred clusters correspond to functionally distinct tumor and microenvironmental compartments.

### Semantic-gated fusion is necessary for GreS performance

After demonstrating the empirical performance of GreS across multiple benchmarks, we investigated whether the incorporation of gene-level semantic information is responsible for the observed performance gains. To this end, we conducted an ablation study on the DLPFC dataset to directly evaluate the contribution of the semantic branch and the semantic-gated fusion mechanism.

We compared the original GreS model with three variants that selectively disrupt semantic information: (i) w/o Semantic, where the semantic branch is removed and the model relies solely on gene expression and spatial graphs; (ii) Shuffle, where semantic embeddings are randomly permuted across spots, preserving their global distribution but destroying spot-level semantic correspondence; and (iii) Random, where semantic embeddings are replaced with randomly initialized vectors. Detailed ARI results across 12 DLPFC sections are reported in Table 4.

**Table 4.** Detailed ARI scores (scaled by 100) of the ablation study on the DLPFC dataset. GreS denotes the original model. w/o Semantic removes the semantic branch. Shuffle permutes semantic embeddings across spots. Random uses randomly initialized embeddings. The best performance is highlighted in **bold**, and the second best is underlined.

| Slice | GreS | w/o Semantic | Shuffle | Random |
| --- | --- | --- | --- | --- |
| 151507 | **75.41** | 67.96 | <u>69.94</u> | 68.88 |
| 151508 | **68.19** | 61.36 | <u>66.70</u> | 57.16 |
| 151509 | **78.40** | 65.52 | <u>69.47</u> | 67.85 |
| 151510 | **74.71** | 65.93 | <u>71.93</u> | 71.81 |
| 151669 | **67.90** | 52.92 | 50.14 | <u>53.76</u> |
| 151670 | **65.96** | 50.38 | 49.34 | <u>51.36</u> |
| 151671 | **85.98** | <u>73.26</u> | 70.16 | 68.81 |
| 151672 | **85.20** | 78.97 | 79.30 | <u>80.53</u> |
| 151673 | **57.82** | 54.34 | <u>55.03</u> | 54.42 |
| 151674 | **55.21** | 50.02 | 51.66 | <u>52.15</u> |
| 151675 | **53.07** | <u>50.52</u> | 51.12 | 46.22 |
| 151676 | **58.41** | 42.98 | <u>44.16</u> | 43.69 |
| **Avg.** | **68.86** | 59.51 | <u>60.74</u> | 59.73 |

Across all sections, the original GreS model achieved the highest performance, with an average ARI of 0.6886. In contrast, all variants that removed or disrupted semantic information exhibited substantially lower performance, with average ARI values ranging from 0.5951 to 0.6074, representing a clear gap relative to the original model. Notably, the three semantic-disrupted variants (w/o Semantic, Shuffle, and Random) achieved comparable performance levels, and none consistently outperformed the others across sections. This pattern indicates that the performance gains of GreS cannot be attributed to the mere presence of additional embeddings or random semantic signals. Instead, effective improvement arises specifically from gene-level semantic information that is biologically meaningful and properly aligned at the spot level.

Together, these results provide direct evidence that gene-level semantic information, when integrated through semantic-gated fusion, is a key contributor to the effectiveness of GreS, rather than an incidental byproduct of increased model complexity or additional feature inputs.

## Discussion

In this paper, we introduce GreS, a spatial domain identification framework that incorporates gene-level semantic priors into spatial representation learning. We evaluated GreS across layered brain tissue, embryonic development, and heterogeneous tumor microenvironments, where it consistently produced spatial domains that are structurally coherent and biologically interpretable. These results indicate that gene-level semantic information can serve as a broadly applicable and complementary source of prior knowledge for spatial modeling, particularly in datasets where expression measurements are sparse or difficult to interpret in isolation.

Recent advances in spatial domain identification have increasingly emphasized the importance of neighborhood information beyond simple physical adjacency. In particular, many methods seek to enrich local context by incorporating expression similarity or dynamically defined pseudo-neighbors, motivated by the observation that spatial proximity alone may be insufficient to characterize boundary or transitional spots. At a conceptual level, these approaches share a common goal, providing each spot with more informative context to support representation learning.

GreS builds on this line of thinking by introducing gene-level semantic information as a means of characterizing the functional state of each spot and guiding how neighborhood information is utilized. Rather than redefining neighborhood graphs, GreS explicitly models two complementary views—a spatial graph capturing physical adjacency and a feature graph capturing expression-driven similarity—and uses semantic priors to modulate their relative contributions. By aggregating functional context across multiple genes, semantic representations provide an informative signal for determining whether spatial continuity or expression similarity is more relevant for a given spot. In this sense, semantic priors act as an inductive bias that helps resolve ambiguity when spatial or expression cues alone are insufficient.

Despite its effectiveness, GreS relies on the availability of gene functional annotations and regulatory network information to construct semantic priors. In biological systems or organisms where such knowledge is incomplete, semantic representations may be less informative. Addressing this dependency will require either continued expansion of biological knowledge resources or the development of strategies that can infer or approximate semantic context when curated information is limited.

Overall, GreS demonstrates that gene-level semantic information can be productively integrated into spatial domain identification as a guiding prior, refining how spatial and expression-based signals are interpreted. More broadly, this work supports a knowledge-informed direction for spatial transcriptomics analysis, in which curated biological information complements data-driven modeling to improve robustness and interpretability in complex tissues.

## Author contributions statement

Y.X, Y.L, J.F, and Y.Z conceived the study. Y.X, Y.L performed analysis. Y.Z. supervised the project. Y.X, Y.L, J.F and Y.Z. wrote the article. All authors read and approved the final article.

## Acknowledgments

This work is supported by the National Natural Science Foundation of China (No. 32500550). Y.Z is partially supported by the Ministry of Human Resources and Social Security of the People’s Republic of China (No. Y20250128), and Guangdong Science and Technology Department. Y.X, and Y.L is support by the Red Bird MPhil (RBM) program at the Hong Kong University of Science and Technology (Guangzhou).

