## Supplementary Table~S1 for "Semantic-Guided Spatial Representation Learning for Spatial Domain Identification"

Table S1: Summary of spatially resolved transcriptomics datasets.

| Platform | Spatial resolution | Tissue | Slice | Spots | Genes | Clusters |
|---|---|---|---|---|---|---|
| 10X Visium | 55 $\mu$m | Human dorsolateral prefrontal cortex (DLPFC) | 151507 | 4226 | 33538 | 7 |
| | | | 151508 | 4384 | 33538 | 7 |
| | | | 151509 | 4789 | 33538 | 7 |
| | | | 151510 | 4634 | 33538 | 7 |
| | | | 151669 | 3661 | 33538 | 5 |
| | | | 151670 | 3498 | 33538 | 5 |
| | | | 151671 | 4110 | 33538 | 5 |
| | | | 151672 | 4015 | 33538 | 5 |
| | | | 151673 | 3639 | 33538 | 7 |
| | | | 151674 | 3673 | 33538 | 7 |
| | | | 151675 | 3592 | 33538 | 7 |
| | | | 151676 | 3460 | 33538 | 7 |
| | | Human breast cancer (HBC) | Section 1 | 3798 | 36601 | 20 |
| Stereo-seq | ∼0.5 $\mu$m | Mouse embryo (ME) | E9.5_E1S1 | 5913 | 25568 | 12 |
| | | | E9.5_E2S1 | 5292 | 23756 | 14 |
| | | | E9.5_E2S2 | 4356 | 24107 | 13 |
| | | | E9.5_E2S3 | 5059 | 24238 | 13 |
| | | | E9.5_E2S4 | 5797 | 23398 | 13 |
